# A robust balancing mechanism for spiking neural networks

**DOI:** 10.1101/2023.08.28.555064

**Authors:** Antonio Politi, Alessandro Torcini

## Abstract

Dynamical balance of excitation and inhibition is usually invoked to explain the irregular low firing activity observed in the cortex. We propose a robust nonlinear balancing mechanism for a random network of spiking neurons, in absence of strong external currents. The mechanism exploits the plasticity of excitatory-excitatory synapses induced by short-term depression. A simple self-consistent analysis accompanied by direct simulations shows the emergence and stability of a balanced asynchronous state in the thermodynamic limit. This regime is essentially fluctuation driven and characterized by highly irregular spiking dynamics of all neurons.

---

The behavior of the mammalian brain follows from a complex interplay between microscopic (single neuron) and macroscopic features. The brain is typically viewed as an oscillator network, and many dynamical regimes can be interpreted making reference to the different forms of synchronization spontaneously emerging in such networks [1–3]. In fact, pulse-coupled phase oscillators, purposely introduced to describe neuronal dynamics, reproduce a large variety of phenomena [4–9].

In mean-field models of globally coupled oscillators, the stationary regime is often found to be asynchronous, i.e. characterized by constant collective features (such as the local field potential) possibly accompanied by tiny fluctuations resulting from the finiteness of the neuronal population. Partial synchrony may manifest itself as either periodic macroscopic oscillations [5, 10], or irregular fluctuations [11, 12]. Anyway, the corresponding single-neuron firing activity is typically regular (the coefficient of variation (CV) of the interspike intervals is small), contrasting the experimental evidence that cortical neurons *in vivo* operate erratically and with a relatively low firing rate [13] in spite of receiving stimulations from thousands of pre-synaptic neurons [14–16].

An irregular firing activity is generated if the neurons operate in the so-called fluctuation-driven regime, when they stay in proximity of the firing threshold, crossed at random times thanks to self-generated fluctuations [17]. This can happen when inhibition is strong and accompanied by a random connectivity which suppresses coherence across the neuronal population [18].

Altogether, it is widely accepted that the underlying dynamics is the so-called *balanced* regime [19], observed in networks characterized by an overall strong synaptic coupling, resulting from the linear superposition of many contributions – an assumption consistent with optogenetic experiments *in vitro* [20]. A balanced state can be, e.g., found, by assuming: (i) a sufficiently large in-degree *K*; (ii) coupling strengths of order 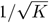; (iii) external currents of order 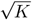 [19, 21–28]. If the external currents are of order 𝒪(1), excitation and inhibition still balance each other, but the firing activity decreases as 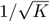, suggesting that strong external currents are a necessary ingredient. However, this latter hypothesis has recently received several criticisms [29, 30] based on experimental evidence that the external input is *𝒪*(1) [31–33].

In this Letter, we show that the introduction of nonlinearities can robustly sustain and stabilize a balanced regime, where the irregular firing of excitatory and inhibitory neurons compensate each other, without neither the inclusion of strong external currents, nor the *ad hoc* adjustment of parameter values. The nonlinear mechanism, herein invoked is the well known short-term synaptic depression (STD) [34], arising from the finitude of available resources [35]. It has been shown that depression has a prevalent effect on excitatory synapses in the visual cortex, inducing dynamical variations of the balance between excitation and inhibition [36].

More precisely, we consider a plastic network of pulse-coupled phase-oscillators, where STD modifies nonlinearly the synaptic inputs. For the sake of simplicity and consistently with experimental indications [36, 37], STD is assumed to act only on the synapses connecting excitatory neurons. We show that this little adjustment suffices to ensure the self-sustainement of an irregular activity.

## The model

As shown in [4], pulse-coupled spiking neurons can be equivalenty modelled by either using the membrane potential as the relevant variable, or by introducing a phase-like variable *ϕ* such that its evolution in the absence of coupling is perfectly linear in time. In the latter case, the specificity of the neuron is encoded in the phase response curve (PRC) [1, 38]. Here, we adopt the second representation as it offers the possibility to compare with a large class of models. More precisely, we consider two neural populations, described by the phase-like variable 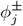, where the upper index “*±*” identifies their excitatory/inhibitory nature. When 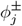 reaches the threshold 1, it is reset to 0, and simultaneously a smooth post-synaptic *α*-pulse *p*_*α*_(*t*) = *α*^2^*t*e^−*αt*^ is delivered to all the connected neurons, mimicking a non-istantaneous synaptic transmission [4, 39, 40]. Mathematically,

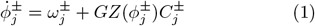

where 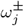 is the bare firing rate, simulating the effect of a “weak” external current due to the projections of neurons outside the recurrent circuit; *G* denotes the global coupling strength; *Z*(*ϕ*) is the above mentioned PRC. The much studied Leaky Integrate-and-Fire (LIF) neurons [41] are characterized by an exponential profile *Z*_LIF_(*ϕ*) = exp(*ϕ* − 1) (normalized to have the maximum equal to 1) [4, 42]. In this Letter we have mostly considered *Z*_I_(*ϕ*) = 12(1 − *ϕ*)*/*[5 +(2 − 2*ϕ*)^6^] (the two shapes are represented in Fig. 1(a)) for its continuity at threshold, as usually assumed in realistic PRCs [43], and its resemblence to PRC for type I membrane excitability [44]. Anyway, also the LIF model has been simulated, confirming the roubstness of our results.

**FIG. 1.**
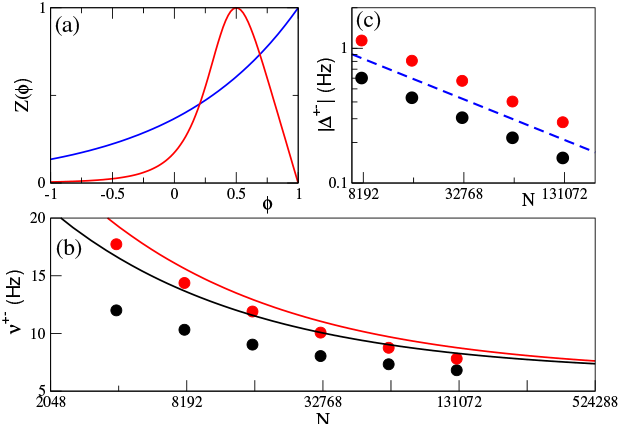
(a) The PRC *Z*(*ϕ*) vs the phase-like variable *ϕ* for *Z*_I_ (red curve) and *Z*_LIF_ (blue curve); (b) the population firing rates 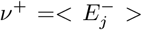 and *ν*^−^ =*< I*_*j*_ *>* versus *N* ; (c) the average unbalance |Δ^*±*^| versus *N*, the blue dashed line denotes a power law decay as 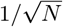. In (b-c) the black (red) color refers to excitatory (inhibitory) neurons and the solid lines in (b) to the self-consistent approximations.

The auxiliary variables 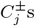 represent the incoming synaptic currents

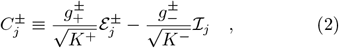

where the 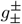 coefficients quantify the specific intra and inter synaptic strengths of excitatory and inhibitory populations, while *K*^*±*^ is the average in-degree, and ℐ_*j*_ and 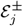 represent the incoming inhibitory and (effective) excitatory fields. The inhibitory field obeys the differential equation

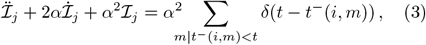

where *α* is the inverse pulse-width, while *t*^−^(*i, m*) denotes the sending time of the *m*-th spike from the *i*-th inhibitory neuron to the *j*-th neuron. This representation amounts to assuming that ℐ_*j*_(*t*) is the linear superposition of the *α*-pulses received by the neuron *j* until time *t*. For large *α*-values, *p*_*α*_ is well approximated by a *δ*-pulse [5]. We prefer to assume *α*-pulses because they are more realistic, accounting in a smooth way for a non-instantaneous transmission [4, 39, 40]. The excitatory field is treated in a slightly different way,

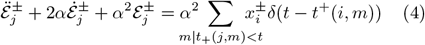

where *i* identifies the excitatory neuron sending the *m*-th spike to the *j*-th neuron; *x*^*±*^ ∈ [0, 1] represents the synaptic efficacy. If the receiving neuron is inhibitory *x*^−^ ≡ 1, while *x*^+^ is affected by the STD acting on excitatory-to-excitatory connections. By following [45], its evolution can be written as

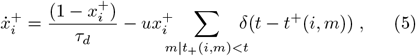

where *t*^+^(*i, m*) identifies the time of the *m*-th spike emitted by the *i*-th neuron itself. Whenever the neuron spikes, the synaptic efficacy 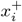 is reduced by a factor *u*, representing the fraction of resources consumed to produce a post-synaptic spike.

So long as as the *i*-th excitatory neuron does not spike, the variable 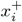 increases towards 1 over a time scale *τ*_*d*_. The network is made of one excitatory and one inhibitory population, each composed of *N* neurons, with the indegrees distributed as in an Erdös-Renyi random graph, massively coupled [46], i.e. with *K*^*±*^ = *c*^*±*^*N* .

## Self-consistent approach

Before discussin the direct numerical simulations, we present a simple self-consistent approach to explain how STD can actually stabilize a balanced state even in absence of strong external currents. In the above defined setup, 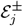 and 𝒪_*j*_, being proportional to the in-degree, are also proportional to *N*, so that the two terms in Eq. (2) are both proportional to 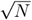. It is useful to make this dependence explicit, by writing

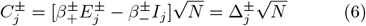

where 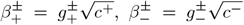. 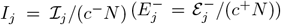 represent the average firing rates of the inhibitory (excitatory) pre-synaptic spike trains stimulating the *j*-th neuron; finally, 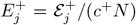 is the effective firing rate of an excitatory neuron scaled to account for the reduced efficacy due to STD. The approximation consists in neglecting neuron-to-neuron fluctuations, as well as temporal variations so that we can drop the *j* dependence of both the fields and the input currents and assume that they are constant. Within this approximation, a balanced regime can exist if *C*^*±*^ remains finite for *N* → ∞, or, equivalently, if the terms in square brackets in Eq. (6) converge to 0 (as 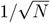). Accordingly,

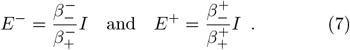

In the absence of STD, *E*^+^ = *E*^−^ and the two conditions can be satified only if 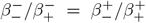 (i.e., 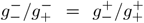): in these conditions the balanced regime is not generic. In literature, a way out has been proposed assuming that the external current *ω*^*±*^ is of order 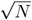 so that it must be included in the balance conditions, thus making the corresponding linear system to solve inhomogeneous [19].

STD, herein included, offers a more general mechanism, making the the homogenous system (7), nonlinear. In fact, *E*^+^ = *x*^+^*E*^−^ where the synaptic efficacy *x*^+^ is itself a function of the (excitatory) field. In the thermodynamic limit (*N* → ∞), the balance condition is achieved whenever the self-determined efficacy assumes the value

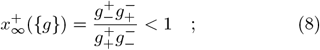

where the inequality follows from the definition of *x*^+^: this is the only required condition to ensure a balanced regime. In practice, we have turned an equality into a much more generic inequality. In this Letter, since we set 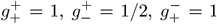, and 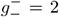, the inequality is satisfied (1*/*2 *<* 1). Superficially, this condition resembles that one found for the balanced regime with strong external currents [19]: in such a context it involves the ratio between excitatory and inhibitory external currents, while here the STD value.

## Mean Firing Rates

The self-consistent analysis is useful to identify the necessary conditions for the onset of a balanced regime, but it can only predict a current-driven regime. In order to analyse the actual network behavior it is necessary to perform numerical simulations. Here below, we report the results for a a homogenous network, where all the bare firing rates are set to 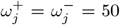 Hz ∀*j* and the PRC is *Z*_*I*_ (*ϕ*) (see [47] for the other parameter values).

In Fig. 1(b) we plot the population firing rates of excitatory 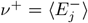 and inhibitory *ν*^−^ = ⟨*I*_*j*_ ⟩ neurons (⟨·⟩ denotes an average over all neurons of a given population) versus *N* . The data are well fitted by the behavior 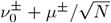, with 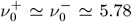 Hz, *μ*^+^ ≃ 399 Hz, and *μ*^−^ *≃* 762 Hz (curves not shown). This indicates that the single-neuron activity remains finite for *N* → ∞, a clear signature of a balanced regime. This conclusion is confirmed by the *N* -dependence of the average unbalance 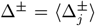, reported in Fig. 1(c), where a clear 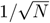 decrease is visible, implying that the average value of *C*^*±*^ stays constant for *N* → ∞.

It is instructive to compare the numerical results with the semi-analytical perturbative implementation of the self-consistent approximation. A detailed account is presented in the supplemental material [48]; here we sketch the main points. Given 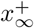 as defined by Eq. (8), one can easily determine the asymptotic ISI *T*_∞_ of the excitatory neurons by integrating Eq. (5) in the presence of a periodic firing activity,

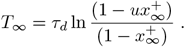

As a result, 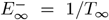 (for our parameter values, 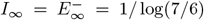 Hz ≃ 6.487 Hz [49]). From the knowledge of the fields, one can determine the input currents 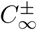 which generate those field, by integrating Eq. (1) under the assumption of a constant current. Finally, the definition (6) of *C*^*±*^ can be used as a consistency relationship to determine the finite-size corrections for both fields, which turn out to be proportional to 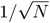. The resulting analytic expressions are reported in [48] and plotted in Fig. 1(b) (see the solid curves). They overestimate the numerical values, but are not too far from them.

## Fluctuations

We start investigating whether the collective dynamics of the network remains asynchronous even for large system sizes, as expected in brain circuits [21]. The raster plot for *N* = 16000, reported in the inset of Fig. 2(a) does not reveal any population oscillation. A more quantitative analysis has been made by computing the time-average of the standard deviation of the incoming fields (i.e. of the instantaneous firing rates), here denoted with *σ*^*±*^. The values computed for different network sizes reported in Fig. 2(a) decrease consistently with the 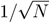 scaling expected from the central limit theorem for an asynchronous dynamics.

**FIG. 2.**
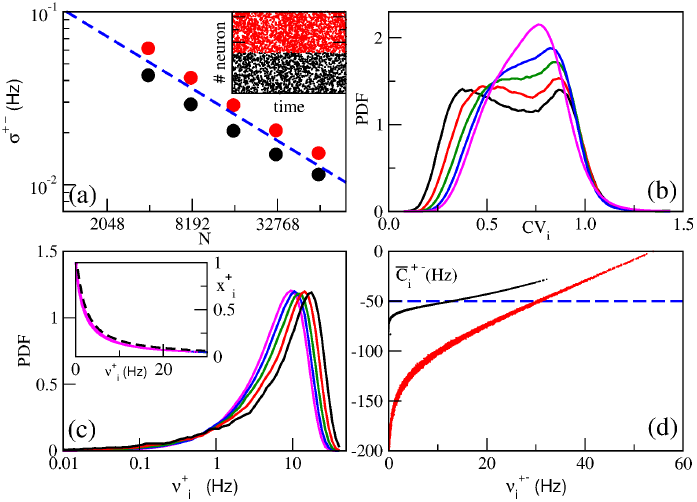
(a) Standard deviation *σ*^*±*^ of the population firing rates versus *N*, the blue dashed line indicates a power law decay as 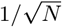. The inset displays a raster plot for *N* = 16000 over a time window of 20 ms. PDF of the coefficients of variation CV_*i*_ (b) and of the firing rates 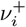 (c) for excitatory neurons. In the inset of (c) is displayed the synaptic efficacy 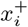 versus the corresponding firing rate 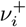, the dashed line is the self-consistent estimation (9). (d) Time averaged values 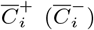 versus the corresponding firing rates 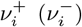 for *N* = 16000. The dashed line denotes the value *C*^*±*^ = −50 Hz discriminating fluctuation from current driven dynamics. The black (red) color in (a-d) refers to excitatory (inhibitory) neurons, while the colors in (b-c) to different system sizes : namely, *N* = 8000 (black); *N* = 16000 (red); *N* = 32000 (green); *N* = 64000 (blue) and *N* = 128000 (magenta).

Next, we focus on temporal fluctuations at the single-neuron level. They are disregarded a priori by the self-consistent approach, but the probability distribution density (PDF) of the coefficient of variations CV_*i*_ reported in Fig. 2(b) for the excitatory neurons gives a clear evidence of irregularity. Some neurons are even characterized by a CV larger than 1, the value expected for a Poisson distribution, and the irregularity tends to increase with *N* . A similar scenario is exhibited by inhibitory neurons (data not shown).

Finally, we turn our attention to ensemble fluctuations. The firing rates themselves are broadly distributed from nearly vanishing values (almost silent neurons) up to 50-60 Hz, with a pronounced peak around 5-10 Hz. When *N* is increased, the PDF widths remain finite and their shapes appear to converge to a given asymptotic form, as clearly visible in Fig. 2(c) where the data refer to excitatory neurons. This manifestation of heterogeneity is not surprising in a massively coupled Erdös-Renyi network. In fact, the single-neuron connectivity is expected to exhibit fluctuations of order 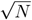, which transform themselves into fluctuations of 𝒪(1) for the 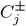, and therefore for the firing rates.

The distribution of firing rates *ν*^+^ induces a distribution of synaptic efficacies *x*^+^ (taken in correspondence of the spiking times). Under the approximation of negligible temporal fluctuations, one can derive the following relationship [48]

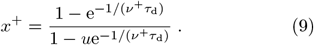

The inset in Fig. 2(c) reveals a good agreement with the numerical simulations.

The PDF shapes reported in Fig. 2(c), are similar to those measured experimentally in the cortex and hippocampus [50–54], with many neurons exhibiting a low firing rate and a high-frequency long tail, akin to a log-normal distribution. These shapes are typically interpreted as an indication of fluctuation-driven dynamics [55]. It is therefore convenient to test whether the neurons, in our model, operate either above or below threshold. This can be done as follows. From Eq. (1), since the maximum of *Z*(*v*^*±*^) is 1, 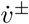 may have a stable zero, only if *ω*^*±*^ + *GC*^*±*^ *<* 0. Hence, a neuron characterized by an average current 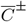 is on average below threshold if 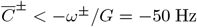. The data reported in Fig. 2(d) reveal a mixed behavior: depending on their effective firing rate, neurons may be either fluctuation- or current-driven. By further averaging over the entire population, we find that while the inhibitory neurons are significantly fluctuation driven with 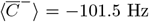, on average the excitatory neurons operate slightly below threshold being 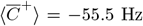. Altogether, the network stabilizes itself in a regime, where fluctuations play a major role, consistently with the observation of a pseudo log-normal distribution of the firing rates [55].

## Robustness of the mechanism

Additional simulations performed for different parameter values and introducing heterogeneity in the external currents have shown the generality of the mechanism. Details can be found in [48]. Here we focus on the most important test, made by using the PRC of LIF neurons, *Z*_LIF_. All parameters have been left unchanged, except for a faster synaptic transmission *α* [47] to check the specificity of the pulsewidth. In Fig. 3 (a) we report the firing rates of the two populations (black and red dots), together with the outcome of the self-consistent approach (solid curves). The theoretical curves converge, as they should, to the same value 1*/T*_∞_, which coincides with the previous asymptotic value since, from its definition, it does not depend on the PRC shape. Also the numerically determined firing rates converge to the same value, again following the law 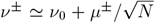, where *ν*_0_ ≃ 5.72 Hz, *μ*^+^ ≃ 480 Hz and, and *μ*^−^ = 803 Hz (curves not shown). The smallness of the discrepancy with the previous asymptotic value (1%) indicates that the irrelevance of the PRC extends to the complete model, where all kinds of fluctuations are automatically included.

**FIG. 3.**
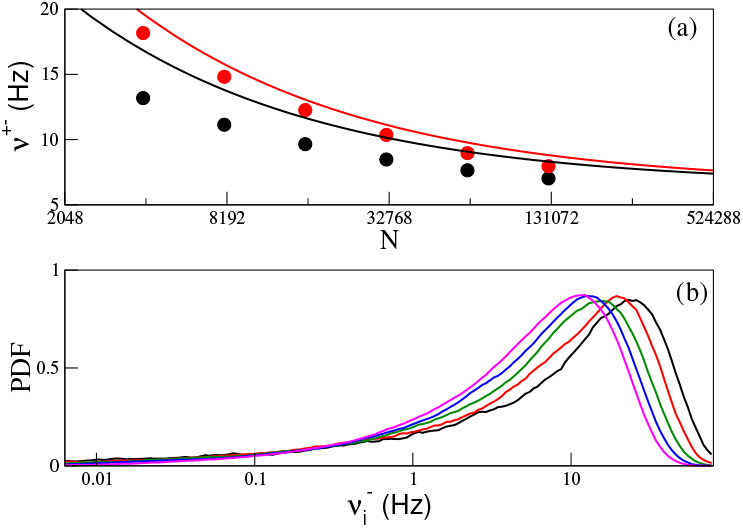
(a) The average population firing rates *ν*^+^ (black circles) and *ν*^−^ (red circles) versus *N* ; (b) PDF of the inhibitory firing rates 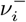. In (a) the black (red) solid line indicates the self-consistent approximation for excitatory (inhibitory) neurons and the colors in (b) different system sizes coded as in Fig. 2 (b-c). The results here reported refer to the LIF model.

In Fig. 3 (b), we plot the PDF of the inhibitory-neuron firing rates for the different network sizes. All curves display a long tail towards vanishing rates, a typical feature of neurons which operate below threshold. Further details of the LIF network dynamics are presented in [48]; they all confirm the typical scenario of a balanced regime as for the *Z*_I_(*ϕ*) PRC.

## Conclusions

The typical setup studied in the literature to discuss dynamically balanced regimes requires the presence of strong external inputs [19]. An alternative layout, which does not suffer this limitation, was proposed in Ref. [30], together with the concept of *sparse balance*. It, however, requires an anomalously broad distribution of synaptic strengths and leads to a vanishing fraction of active neurons (in the thermodynamic limit). The mechanism proposed here is more general and robust: it exploits the dynamical adjustment of the synaptic currents induced by nonlinearities. The non-linear mechanism discussed in this Letter, is short-term synaptic depression (STD), which, lowering the strength of highly active excitatory connections, binds the activity of excitatory neurons. STD is a much studied mechanism, already invoked to explain fundamental cognitive functions, such as working memory [56–58] and the internal representation of spatio-temporal information [59–62]. We expect that other nonlinear mechanisms can be generically effective in ensuring a balanced regime in absence of strong inputs. Spike-frequency adaptation is a mechanism that would be worth exploring since it has been already shown to ensure balance in networks with highly heterogenous in-degrees [63], although still in a setup with strong external currents. The same can be said of facilitation, found to promote the emergence of bistable balanced regimes [64], again in the presence of strong external currents.

We acknowledge useful discussions with German Mato on the initial stage of development of this project during our stay at Max Planck Institute for the Physics of Complex Systems (Dresden, Germany) within the Advanced Study Group “From Microscopic to Collective Dynamics in Neural Circuits” (2016/17). A. T. received financial support by the ANR Project ERMUNDY (Grant No. ANR-18-CE37-0014), by the Labex MME-DII (No. ANR-11-LBX-0023-01) and by CY Generations (Grant No ANR-21-EXES-0008), all part of the French programme “Investissements d’Avenir”. A.P. received support by CY Advanced Studies (Cergy-Pontoise, France) for a visiting scholarship in 2018.

## References

[1] A. T. Winfree, The Geometry of Biological Time, 2nd ed., Interdisciplinary Applied Mathematics, Vol. 12 (Springer-Verlag New York, 2001).

[2] S. H. Strogatz and I. Stewart, Scientific american 269, 102 (1993).

[3] A. Pikovsky and M. Rosenblum, Chaos: An Interdisciplinary Journal of Nonlinear Science 25, 097616 (2015).

[4] L. F. Abbott and C. Van Vreeswijk, Physical Review E 48, 1483 (1993).

[5] C. van Vreeswijk, Physical Review E 54, 5522 (1996).

[6] M. Timme, F. Wolf, and T. Geisel, Physical review letters 89, 258701 (2002).

[7] T. P. Vogels, K. Rajan, and L. F. Abbott, Annu. Rev. Neurosci. 28, 357 (2005).

[8] H. Haken, Brain dynamics: synchronization and activity patterns in pulse-coupled neural nets with delays and noise (Springer Science & Business Media, 2006).

[9] S. Olmi, A. Politi, and A. Torcini, Europhysics Letters 92, 60007 (2011).

[10] X.-J. Wang and G. Buzsáki, Journal of neuroscience 16, 6402 (1996).

[11] S. Luccioli and A. Politi, Phys. Rev. Lett. 105, 158104 (2010).

[12] E. Ullner and A. Politi, Physical Review X 6, 011015 (2016).

[13] W. R. Softky and C. Koch, Journal of neuroscience 13, 334 (1993).

[14] A. Destexhe and D. Paré, Journal of neurophysiology 81, 1531 (1999).

[15] R. M. Bruno and B. Sakmann, Science 312, 1622 (2006).

[16] S. Lefort, C. Tomm, J.-C. F. Sarria, and C. C. Petersen, Neuron 61, 301 (2009).

[17] M. N. Shadlen and W. T. Newsome, Current opinion in neurobiology 4, 569 (1994).

[18] N. Brunel and V. Hakim, Neural computation 11, 1621 (1999).

[19] C. van Vreeswijk and H. Sompolinsky, Science 274, 1724 (1996).

[20] J. Barral and A. D. Reyes, Nature neuroscience 19, 1690 (2016).

[21] A. Renart, J. de la Rocha, P. Bartho, L. Hollender, N. Parga, A. Reyes, and K. D. Harris, Science 327, 587 (2010).

[22] M. Monteforte and F. Wolf, Phys. Rev. Lett. 105, 268104 (2010).

[23] A. Litwin-Kumar and B. Doiron, Nat Neurosci 15, 1498 (2012).

[24] J. Kadmon and H. Sompolinsky, Phys. Rev. X 5, 041030 (2015).

[25] R. Rosenbaum and B. Doiron, Physical Review X 4, 021039 (2014).

[26] R. Pyle and R. Rosenbaum, Physical Review E 93, 040302 (2016).

[27] M. di Volo and A. Torcini, Phys. Rev. Lett. 121, 128301 (2018).

[28] M. Di Volo, M. Segneri, D. S. Goldobin, A. Politi, and A. Torcini, Chaos: An Interdisciplinary Journal of Nonlinear Science 32, 023120 (2022).

[29] Y. Ahmadian and K. D. Miller, Neuron 109, 3373 (2021).

[30] R. Khajeh, F. Fumarola, and L. Abbott, PLOS Computational Biology 18, e1008836 (2022).

[31] S. Chung and D. Ferster, Neuron 20, 1177 (1998).

[32] I. M. Finn, N. J. Priebe, and D. Ferster, Neuron 54, 137 (2007).

[33] A. D. Lien and M. Scanziani, Nature neuroscience 16, 1315 (2013).

[34] M. Tsodyks and S. Wu, Scholarpedia 8, 3153 (2013).

[35] M. V. Tsodyks and H. Markram, Proceedings of the national academy of sciences 94, 719 (1997).

[36] J. A. Varela, S. Song, G. G. Turrigiano, and S. B. Nelson, Journal of Neuroscience 19, 4293 (1999).

[37] J. A. Varela, K. Sen, J. Gibson, J. Fost, L. Abbott, and S. B. Nelson, Journal of Neuroscience 17, 7926 (1997).

[38] C. C. Canavier, Scholarpedia 1, 1332 (2006).

[39] C. Van Vreeswijk, L. Abbott, and G. B. Ermentrout, Journal of computational neuroscience 1, 313 (1994).

[40] S. Coombes, G. J. Lord, and M. R. Owen, Physica D: Nonlinear Phenomena 178, 219 (2003).

[41] A. N. Burkitt, Biological cybernetics 95, 1 (2006).

[42] A. Politi and M. Rosenblum, Phys. Rev. E 91, 042916 (2015).

[43] R. M. Smeal, G. B. Ermentrout, and J. A. White, Philosophical Transactions of the Royal Society B: Biological Sciences 365, 2407 (2010).

[44] B. Ermentrout, Neural computation 8, 979 (1996).

[45] M. Tsodyks, K. Pawelzik, and H. Markram, Neural computation 10, 821 (1998).

[46] D. Golomb, D. Hansel, and G. Mato, in Handbook of Biological Physics 4, 887 (2001).

[47] The parameters are set as follows. Each neuron has a probability of 10% to be connected to any other neuron, to guarantee that 80% (20%) of these connections are excitatory (inhibitory) as in the cortex we set *c*^+^ = 0.08 (*c*^−^ = 0.02). For the STD parameters we set u = 0.5 (a single spike emission halves the synaptic resources) and *τ*_*d*_ = 1 s. The overall coupling strength has been fixed to *G* = 1. For the *Z*_I_(*ϕ*) (*Z*_LIF_(*ϕ*)) PRC we considered synapses with *α*^−1^ = 0.2 ms (*α*^−1^ = 0.04 ms).

[48] See Supplemental Material at [URL will be inserted by publisher] for details on the self-consistent analysis, for the results on heterogenous networks and on further possible dynamical regimes, as well as on further numerical investigations of the LIF network.

[49] In our case, the inhibitory field is equal to the excitatory field.

[50] T. Hromádka, M. R. DeWeese, and A. M. Zador, PLoS biology 6, e16 (2008).

[51] D. H. O’Connor, S. P. Peron, D. Huber, and K. Svoboda, Neuron 67, 1048 (2010).

[52] A. Wohrer, M. D. Humphries, and C. K. Machens, Progress in neurobiology 103, 156 (2013).

[53] G. Buzsáki and K. Mizuseki, Nature Reviews Neuroscience 15, 264 (2014).

[54] G. Mongillo, S. Rumpel, and Y. Loewenstein, Nature neuroscience 21, 1463 (2018).

[55] A. Roxin, N. Brunel, D. Hansel, G. Mongillo, and C. van Vreeswijk, Journal of Neuroscience 31, 16217 (2011).

[56] P. Del Giudice, S. Fusi, and M. Mattia, Journal of Physiology-Paris 97, 659 (2003).

[57] G. Mongillo, O. Barak, and M. Tsodyks, Science 319, 1543 (2008).

[58] H. Taher, A. Torcini, and S. Olmi, PLOS Computational Biology 16, e1008533 (2020).

[59] S. Romani and M. Tsodyks, Hippocampus 25, 94 (2015).

[60] Y. Wang, S. Romani, B. Lustig, A. Leonardo, and E. Pastalkova, Nature neuroscience 18, 282 (2015).

[61] C. Haimerl, D. Angulo-Garcia, V. Villette, S. Reichinnek, A. Torcini, R. Cossart, and A. Malvache, Proceedings of the National Academy of Sciences 116, 7477 (2019).

[62] B. Pietras, V. Schmutz, and T. Schwalger, PLOS Computational Biology 18, e1010809 (2022).

[63] I. D. Landau, R. Egger, V. J. Dercksen, M. Oberlaender, and H. Sompolinsky, Neuron 92, 1106 (2016).

[64] D. Hansel and G. Mato, Journal of Neuroscience 33, 133 (2013).

